# Sustained ability of a natural microbial community to remove nitrate from groundwater

**DOI:** 10.1101/2021.05.27.446013

**Authors:** Charles J. Paradis, John I. Miller, Ji-Won Moon, Sarah J. Spencer, Lauren M. Lui, Joy D. Van Nostrand, Daliang Ning, Andrew D. Steen, Larry D. McKay, Adam P. Arkin, Jizhong Zhou, Eric J. Alm, Terry C. Hazen

## Abstract

Microbial-mediated nitrate removal from groundwater is widely recognized as the predominant mechanism for nitrate attenuation in contaminated aquifers and is largely dependent on the presence of a carbon-bearing electron donor. The repeated exposure of a natural microbial community to an electron donor can result in the sustained ability of the community to remove nitrate; this phenomenon has been clearly demonstrated at the laboratory scale. However, *in situ* demonstrations of this ability are lacking. For this study, ethanol (electron donor) was repeatedly injected into a groundwater well (treatment) for six consecutive weeks to establish the sustained ability of a microbial community to remove nitrate. A second well (control) located up-gradient was not injected with ethanol during this time. The treatment well demonstrated strong evidence of sustained ability as evident by concomitant ethanol and nitrate removal and subsequent sulfate removal upon consecutive exposures. Both wells were then monitored for six additional weeks under natural (no injection) conditions. During the final week, ethanol was injected into both treatment and control wells. The treatment well demonstrated sustained ability as evident by concomitant ethanol and nitrate removal whereas the control did not. Surprisingly, the treatment well did not indicate a sustained and selective enrichment of a microbial community. These results suggested that the predominant mechanism(s) of sustained ability likely exist at the enzymatic- and/or genetic-levels. The results of this study demonstrated that the *in situ* ability of a microbial community to remove nitrate can be sustained in the prolonged absence of an electron donor. Moreover, these results implied that the electron-donor exposure history of nitrate-contaminated groundwater can play an important role nitrate attenuation.

**Article Impact Statement:** Groundwater microbes sustain ability to remove nitrate in absence of carbon and energy source.

## 1. Introduction

Natural microbial communities that can utilize nitrate as an electron acceptor are ubiquitous in groundwater and play a critical role in nitrate attenuation in contaminated aquifers (Rivett et al. 2008). The ability of these communities to reduce and effectively remove nitrate from groundwater is primarily limited by the availability of a suitable electron donor (Rivett et al. 2008). Prior exposure of a community to an electron donor can result in the sustained ability of the community to conduct specific donor-acceptor reactions (Leahy and Colwell 1990; Kline et al. 2011). This phenomenon has been observed in the field based on characterization studies and has been demonstrated in the laboratory based on experimental studies (Koskella and Vos 2015).

For example, in the field, Pernthaler and Pernthaler (2005) observed the sustained ability of a marine microbial community in response to naturally fluctuating electron donor availability over the course of a single day. In the laboratory, Pernthaler et al. (2001) demonstrated that the sustained ability of marine isolates was dependent on the frequency of electron donor addition, e.g., one species out-competed the other during a single addition whereas the other species performed best during hourly additions. Leahy and Colwell (1990) summarized the predominant, yet inter-related, mechanisms by which sustained ability can occur: (1) induction and/or depression of specific enzymes, (2) genetic changes that result in new metabolic capabilities, and (3) selective enrichment of microbes able to conduct the donor-acceptor reactions of interest. More recently and in the laboratory, Oh et al. (2013) demonstrated the inter-related mechanisms of the sustained ability of a river sediment microbial community to utilize nitrate as an electron acceptor in response to exposures of an electron donor (benzalkonium chlorides); this resulted in both the selective enrichment of *Pseudomonas* species and genetic changes via benzalkonium chlorides-related amino acid substitutions and horizontal gene transfer.

These observations, demonstrations, and mechanistic insights of the sustained ability of natural microbial communities conduct specific donor-acceptor reactions are only a small fraction of those in the vast literature (Koskella and Vos 2015) yet they clearly illustrate the importance and highlight the current understanding of the topic. Nevertheless, there is a need to bridge the knowledge gap between field observations and laboratory demonstrations of sustained ability. Specifically, there is a need to design and conduct highly controlled field experiments with the proper controls to both demonstrate sustained ability and elucidate its mechanisms. The objectives of this study were to: (1) establish a natural microbial community able to utilize nitrate as an electron acceptor in groundwater, (2) determine how long sustained ability can last in the absence of a suitable electron donor, and (3) elucidate the microbial mechanism(s) responsible for sustained ability the community to remove nitrate.

## 2. Materials and Methods

### 2.1. Study site

The study site is in Area 2 of the Y-12 S-3 pond field site which is a part of the Oak Ridge Reservation (ORR) and in Oak Ridge, Tennessee, USA (Fig. 1). The hydrogeology of the study site has been previously described (Paradis et al. 2016; Paradis et al. 2018; Watson et al. 2004). The subsurface consists of approximately 6 meters of unconsolidated and heterogeneous materials comprised of silty and clayey fill underlain by undisturbed and clay-rich weathered bedrock. The study site contains 13 monitoring wells (FW218 through FW230), two of which were used as test wells (FW222 and FW224), and one of which was used as a source well (FW229) for groundwater injectate for the exposure tests, as discussed in Section 2.2. (Fig. 1).

**Fig. 1.**
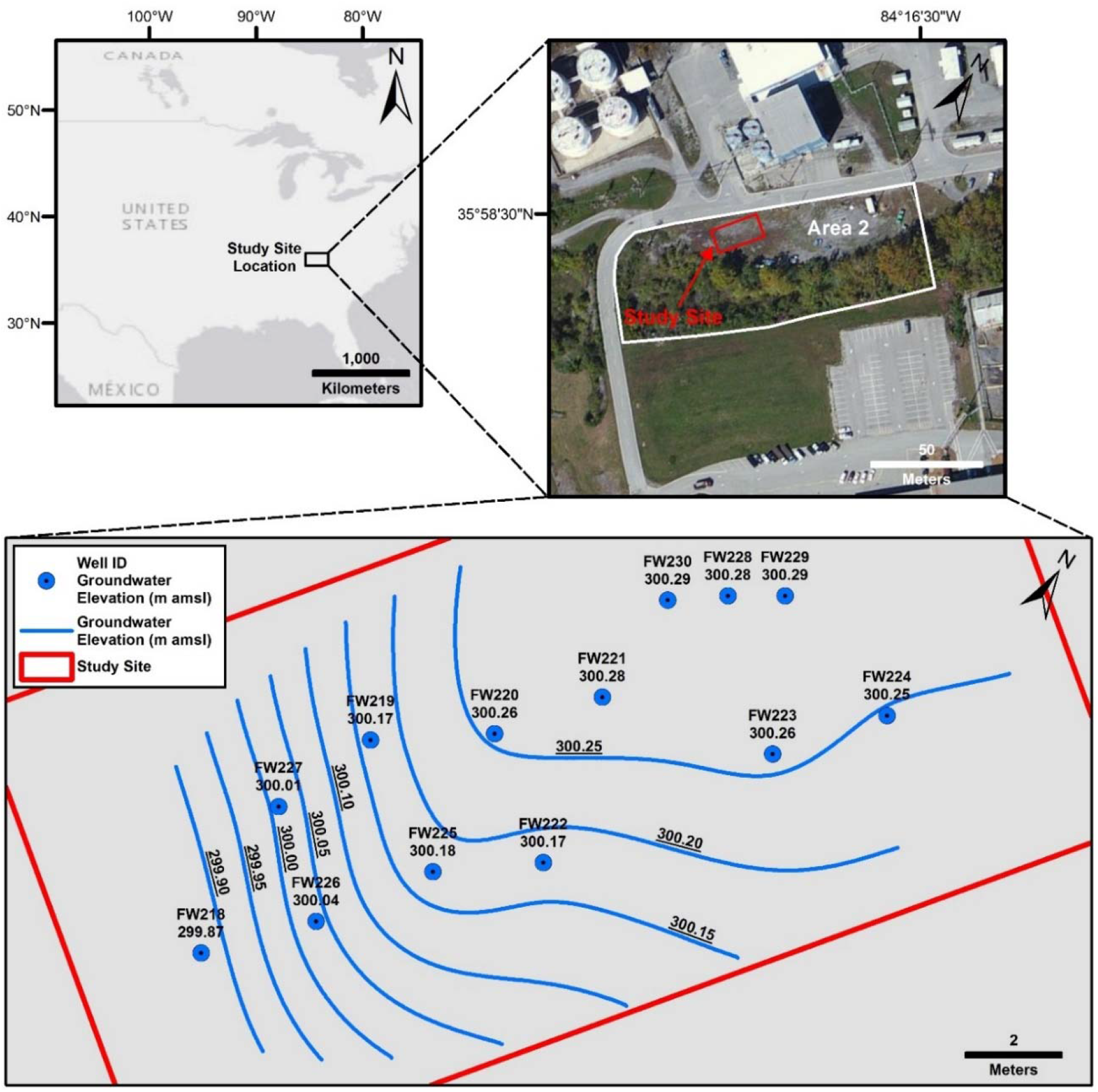
Plan-view maps of the study site from Paradis et al. (2017), clockwise from upper left, country map showing study site location in the southeastern United States, area map showing study site location in Area 2 of the Oak Ridge Reserve, and study site map showing well locations, groundwater elevations, and groundwater elevation iso-contours, m amsl = meters above mean sea level, treatment well is FW222, control well is FW224.

The test wells are constructed of 1.9-cm inside diameter schedule-80 polyvinyl chloride (PVC) pipe and are screened from 3.7 to 6.1 m below ground surface (mbgs). The test wells are screened within the fill materials and were vertically terminated at contact with the undisturbed weathered bedrock. The shallow groundwater aquifer is unconfined and the depth to groundwater is approximately 3.5 mbgs. The groundwater pH is circumneutral (pH ≈ 6.5 to 8.0) and dissolved oxygen (DO) is relatively low (DO ≈ 1 to 2 mg/L). Nitrate and sulfate concentrations range from approximately 5 to 75 and 10 to 200 mg/L, respectively; the groundwater geochemistry has been previously described (Paradis et al. 2016; Paradis et al. 2018; Watson et al. 2004). The test wells are separated by approximately 6 m of horizontal distance and oriented nearly perpendicular to the direction of groundwater flow (Fig. 1).

### 2.2. Electron Donor Exposure Tests

Electron donor exposure tests were conducted using the single-well push-pull test method (Istok 2013). During a push-pull test, a volume of water which contains a known mass of one or more non-reactive and reactive tracers is injected into a single well under forced-flow conditions; this is referred to as the push phase (Fig. 2). The mixture of the injection fluid and aquifer fluid is then collected periodically from the same well under natural-flow conditions; this is referred to as the pull or drift phase (Fig. 2). The concentrations of the added tracers, reactants, and products are then plotted versus the time elapsed to generate breakthrough curves. The breakthrough curves are then analyzed to characterize the mass transport mechanisms within the groundwater system, e.g., advection, dispersion, sorption, and microbial-mediated reactivity.

**Fig. 2.**
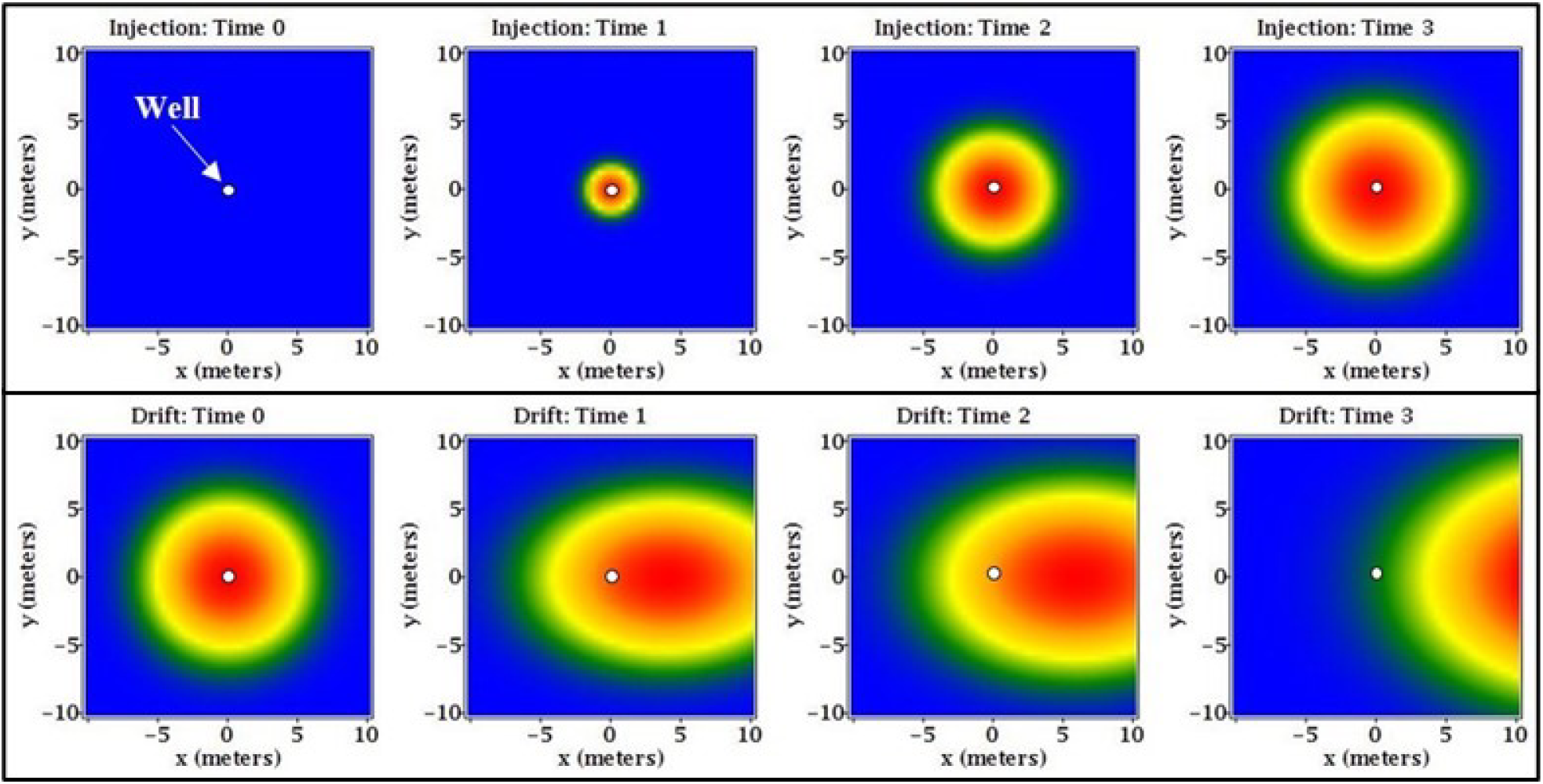
Conceptual model of a single-well push-pull test in plain view showing the forced-flow injection (push) phase (top panel) and the natural-flow drift (pull) phase (bottom panel), blue color represents the aquifer fluid, warmer colors represent the relative concentration of the injection fluid, natural groundwater flow is from left to right.

For this study, a volume of groundwater (5 to 40 L) was collected from up-gradient well FW229 (Fig. 1) using a peristaltic pump and stored in a plastic carboy. A mass of potassium bromide (KBr) (Sigma-Aldrich) and ethanol (C_2_H_6_O) (Sigma-Aldrich) was added to the injection solution and mixed by re-circulation using a peristaltic pump for a target concentration of 200 mg/L bromide and 200 mg/L ethanol. Bromide was added as a non-reactive tracer whereas ethanol was added as a reactive tracer. The addition of ethanol (≈1,400 mg/L) at the study site was previously shown to serve as a suitable electron donor to stimulate nitrate removal (Paradis et al. 2016). The injection solution was then injected into the test well (either treatment or control well), followed by a 20-min resting period, and then periodically sampled over the course of four hours. Immediately prior to, and after mixing of the injection solution, three samples were collected, filtered (0.2 μm filter), stored in 20 mL scintillation vials without headspace, preserved at 4°C, and promptly analyzed for bromide, nitrate, sulfate, and acetate by ion chromatography (Dionex ICS-5000^+^) and for ethanol by gas chromatography (Agilent 6890). Acetate was previously shown to be the predominant metabolite of microbial-mediated oxidation of ethanol under anaerobic conditions from sediments collected within Area 2 at the OR-IFRC (Jin and Roden 2011). Three samples were also collected from the injection well immediately prior to injection and analyzed.

A series of seven exposure tests were conducted in test well FW222 (treatment exposure) and one exposure test was conducted in test well FW224 (control exposure) (Table 1). The treatment was exposed to ethanol for six consecutive weeks (weeks two through seven) followed by six consecutive weeks (weeks eight through thirteen) of no exposure to ethanol (Table 1). During this time, the control was not exposed to ethanol and was subject only to natural hydrogeologic conditions. During week fourteen, both the treatment and control wells were exposed to ethanol (Table 1). The exposure tests allowed for comparing the effects of repeated exposure history (treatment) versus no exposure history (control) in terms of microbial-mediated removal of nitrate.

**Table 1.**
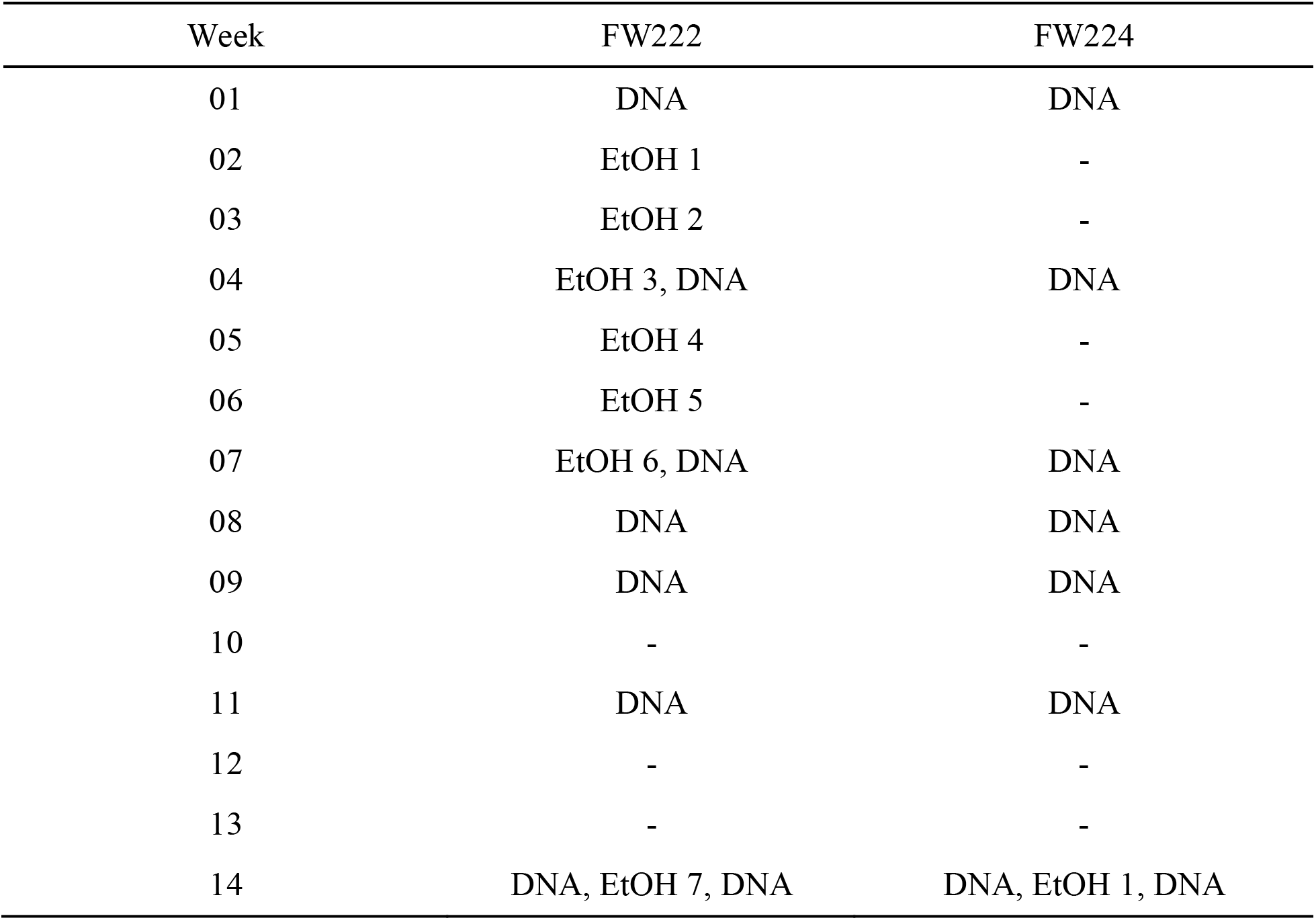
Experimental design of electron donor exposure tests for the treatment well (FW222) and control well (FW224), EtOH = ethanol, DNA = 16S amplicon sequencing of rDNA from planktonic microbes

The breakthrough curves of bromide, ethanol, acetate, nitrate, and sulfate, were analyzed according to the general methodology of Paradis et al. (2019). In brief, three equations were used to characterize natural groundwater flow, non-reactive transport, and reactive transport, respectively, as follows:

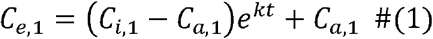

where:

*C*_*e*,1_ = concentration of non-reactive tracer in extraction fluid [L^3^/T]
*C*_*i*,1_ = concentration of non-reactive tracer in injection fluid [L^3^/T]
*C*_*a*,1_ = concentration of non-reactive tracer in aquifer fluid [L^3^/T]
*k* = first-order dilution rate [1/T]
*t* = time elapsed [T]

and

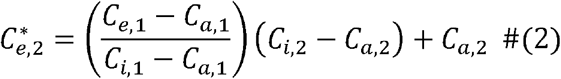

where:

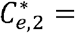 expected concentration of reactive tracer in extraction fluid due to dilution [L^3^/T]
*C*_*i*,2_ = concentration of reactive tracer in injection fluid [L^3^/T]
*C*_*a*,2_ = concentration of non-reactive tracer in aquifer fluid [L^3^/T]

and

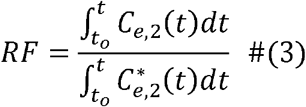

where:

*RF* = recovery factor [dimensionless]
*C*_*e*,2_ = measured concentration of reactive tracer in extraction fluid [L^3^/T]

Equation (1) describes the dilution of the finite volume of injection fluid with respect to the nearly infinite volume of aquifer fluid where the first-order dilution rate (*k*) is proportional to the rate of groundwater flow through the well and its surrounding aquifer material. Equation (2) describes the expected concentration of a reactive tracer in the extraction fluid due to dilution of the injection fluid where any difference between its expected concentration 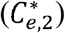 and its measured concentration (*C*_*e*,2_) can be attributed to one or more reactive processes, e.g., microbial-mediated reactivity. Equation (3) describes the ratio of the measured mass recovery of a tracer as compared its expected mass recovery when accounting for dilution. For example, a recovery factor (*RF*) greater than one indicates a net addition of the tracer to the aqueous phase whereas an *RF* less than one indication a net removal of the tracer from the aqueous phase and an *RF* equal to one indicates no change. Equation (3) must be evaluated using numerical integration methods, because the breakthrough curve data is both discrete and its underlying continuous function is unknown. For this study, Equation (3) was evaluated using the mid-point, trapezoid, and Simpson’s techniques and the average *RF* plus or minus its standard error was reported.

### 2.3. Microbial Community Structure

The test wells were sampled for microbial community structure according to the general methodology of Smith et al. (2015). A volume of groundwater (5 to 10 L) was collected from the wells prior to and following the exposure tests. The groundwater was filtered, in series, through a 10 μm and a 0.2 μm filter, and preserved at −80°C. Microbial DNA was extracted from the 0.2 μm filter using a modified Miller method (Hazen et al. 2010; Miller et al. 1999; Smith et al. 2015) and shipped to the Institute for Environmental Genomics (Norman, OK, USA) for analysis of microbial DNA.

Extracted DNA was amplified as described in Wu et al. (2015). DNA was PCR amplified using a two-step PCR. In the first step, 16S rDNA was amplified for 10 cycles using primers 515F and 806R. In the second step, product from the first step was amplified for an additional 20 cycles using primers containing spacers to increase base diversity, barcodes, Illumina adaptor and sequencing primers, and the target primers, 515F and 806R. Amplification efficiency was evaluated by agarose gel electrophoresis. PCR products were pooled in equal molality and purified. Sequencing libraries were prepared according to the MiSeq™ Reagent Kit Preparation Guide (Illumina, San Diego, CA, USA) (Caporaso et al. 2012). Sequencing was performed for 251, 12, and 251 cycles for forward, index, and reverse reads, respectively, on an Illumina MiSeq using a 500-cycle v2 MiSeq reagent cartridge.

The resulting DNA sequences were analyzed according to the general methodology of Techtmann et al. (2015). DNA sequences were analyzed using the QIIME version 1.8.0-dev pipeline (Caporaso et al. 2012) and paired-end raw reads were joined using fastq-join (Aronesty 2015). The joined sequences were demultiplexed and quality filtered in QIIME to remove reads with phred scores below 20. Chimera detection was then performed on joined reads using UCHIME (Edgar 2010; Edgar et al. 2011). Joined, quality-filtered and chimera-checked sequences were deposited at MG-RAST. Sequences were clustered into operational taxonomic units (OTUs, 97% similarity) with UCLUST (Edgar 2010) using the open reference clustering protocol. The resulting representative sequences were aligned using PyNAST (Caporaso et al. 2010) and given a taxonomic assignment using RDP (Wang et al. 2007) retrained with the May 2013 Greengenes release. The resulting OTU table was filtered to keep OTUs that were present at greater than 0.005%, and then rarified to 13,753 sequences per sample (the minimum number of remaining sequences in the samples).

To test the hypothesis that exposure to ethanol influenced community structure, non-metric multi-dimensional scaling (NMDS) and hierarchical clustering analysis (HCA) were performed. A Bray-Curtis dissimilarity matrix was constructed using the scipy.spatial.distance methods from the SciPy library (Jones et al. 2001) in Python (Python 2017) and used as input for NMDS and HCA. NMDS was performed using the sklearn.manifold methods from the Scikit-learn library (Pedregosa et al. 2011). HCA was performed with the scipy.cluster.hierarchy methods using the average linkage method. The number of dimensions was increased starting from two to identify the minimum number of dimensions necessary to achieve a reasonable stress value. A breakpoint was identified at three dimensions, above which ordination stress did not decrease substantially.

## 3. Results and Discussion

### 3.1. Electron Donor Exposure Tests

The breakthrough curves of bromide in the treatment well during the six consecutive weeks of ethanol exposure demonstrated first-order dilution rates (Equation 1) ranging from - 0.69 to −2.16/days (Fig. 3). The dilution rates during the latter three weeks were substantially greater then observed during the first three weeks (Fig. 3). These results indicated that the rate of groundwater flow through the treatment well and its surrounding aquifer material was transient as opposed to steady state. The transient behavior of groundwater flow was not surprising when considering that the aquifer is unconfined and the depth to groundwater is relatively shallow (approximately 3.5 mbgs); these hydrogeologic characteristics make the aquifer particularly sensitive to recharge and discharge events. The breakthrough curves of bromide in the treatment and control wells during the final week of ethanol exposure also demonstrated first-order dilution rates (Fig. 4). However, these rates were relatively low (−0.15 to −0.30/days) as compared to the first six weeks (Fig. 4) and further indicated the transient behavior of groundwater flow. Nevertheless, the rates of groundwater flow during the final week of ethanol exposure in both treatment and control wells were notably similar as evident by dilution rates within a factor of two (Fig. 4). It must be noted that the breakthrough curves bromide (Figs. 3 and 4) were interpreted to represent non-reactive dilution between the injection and aquifer.

**Fig. 3.**
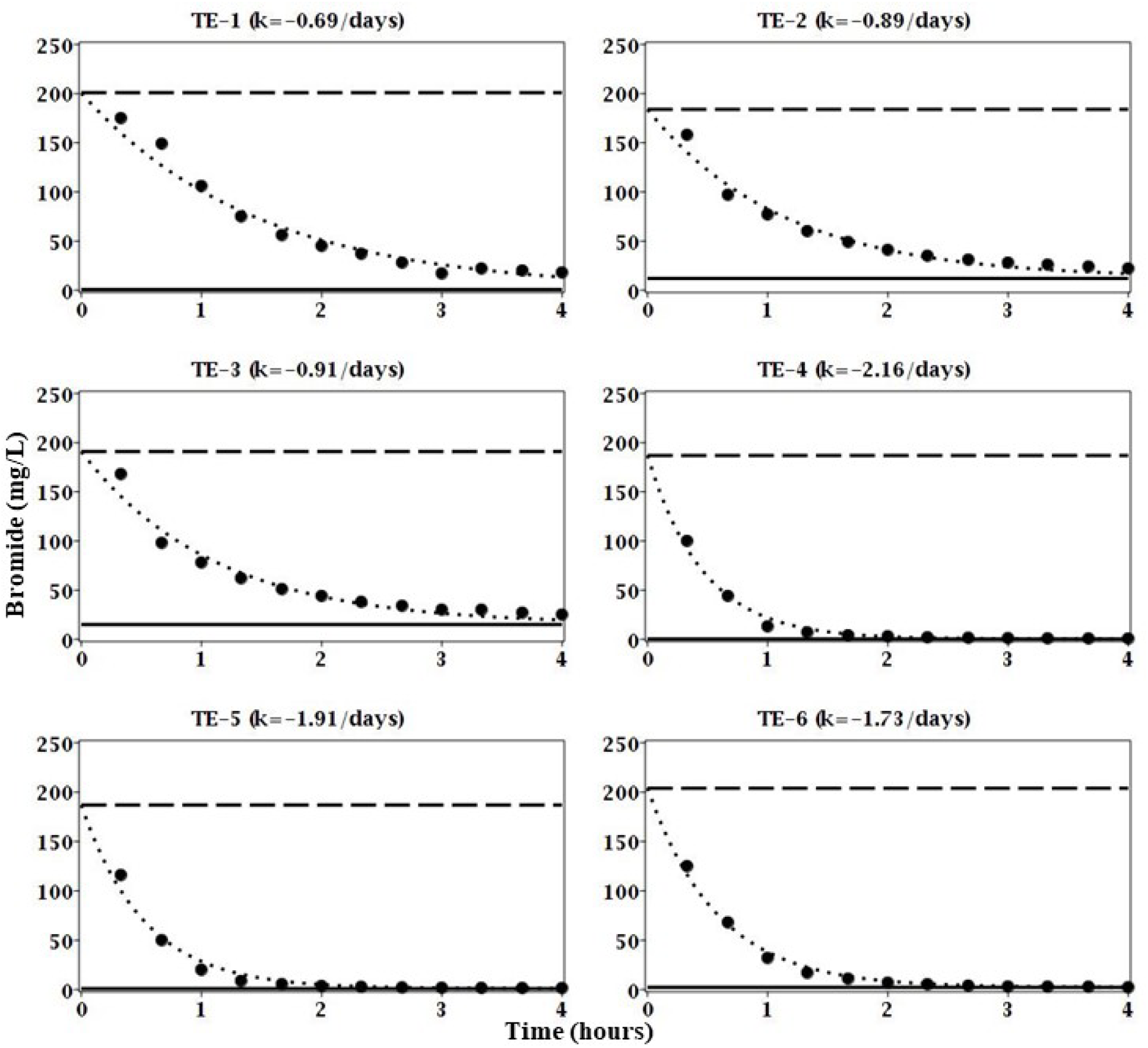
Breakthrough curves of bromide (non-reactive tracer) for treatment exposures 1 through 6 (TE-1 through TE-6) in well FW222), solid circles (●) are concentrations of bromide in the extraction fluid, dashed line (□ □) is the concentration of bromide in the injection fluid, solid line (□ □) is the concentration of bromide in the aquifer fluid or the lower detection limit, dotted line (□ □ □ □) is the best fit of the first-order dilution rate (k).

**Fig. 4.**
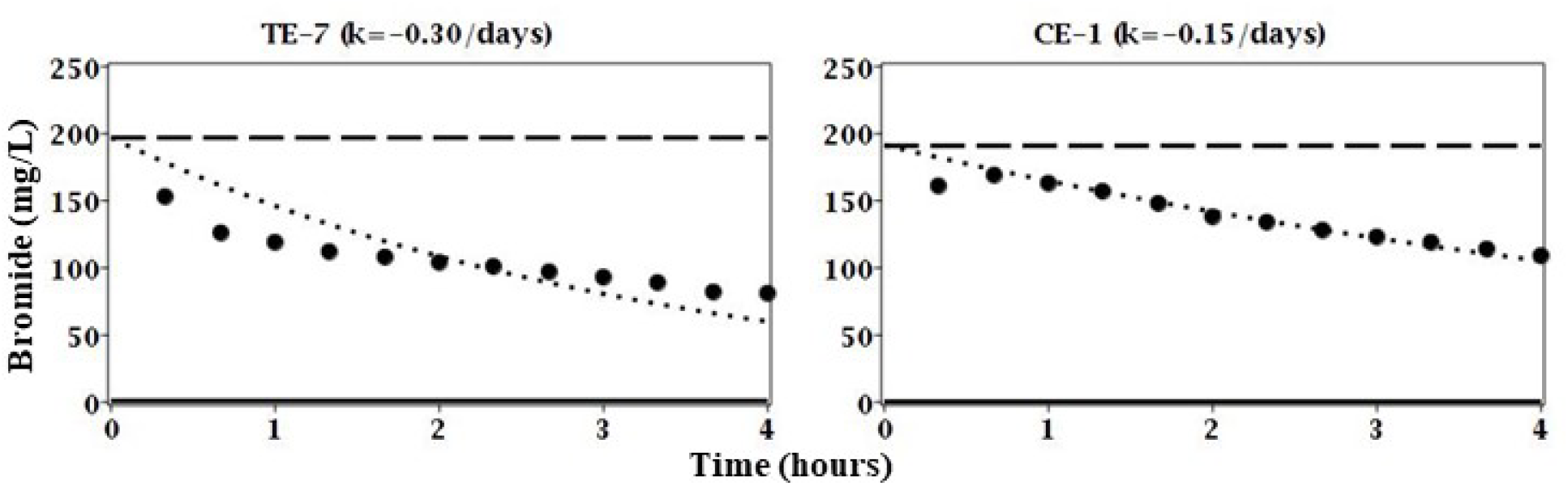
Breakthrough curves of bromide (non-reactive tracer) for treatment exposure 7 (TE-7) well FW222 and control exposure 1 (CE-1) in well FW224, solid circles (●) are concentrations of bromide in the extraction fluid, dashed line (□ □) is the concentration of bromide in the injection fluid, solid line (□ □) is the concentration of bromide in the aquifer fluid or the lower detection limit, dotted line (□ □ □ □) is the best fit of the first-order dilution rate (k).

The breakthrough curves of ethanol, nitrate, and sulfate for exposure one in the treatment well (TE-1) did not demonstrate concomitant removal of ethanol and nitrate or sulfate as evident by the lack of clear and convincing trends in the data or recovery factors (Fig. 5). These results suggested that the natural microbial community was not readily able to utilize ethanol and nitrate. However, the breakthrough curves for exposures two and three (TE-2 and TE-3) did demonstrate concomitant ethanol and nitrate removal and subsequent sulfate removal as evident by substantially lower than expected concentrations; nitrate and sulfate concentrations actually fell below even that of the aquifer fluid (Fig. 5). Microbial-mediated oxidation of ethanol to acetate and reduction of nitrate and sulfate has been well documented at the study site (Wu et al. 2006; Wu et al. 2007) and abroad (Feris et al. 2008; Rodriguez-Escales et al. 2016; Vidal-Gavilan et al. 2014). Moreover, the relative increase in microbial activity during subsequent exposures to ethanol, i.e., sustained ability, was expected based on previous studies (Kline et al. 2011).

**Fig. 5.**
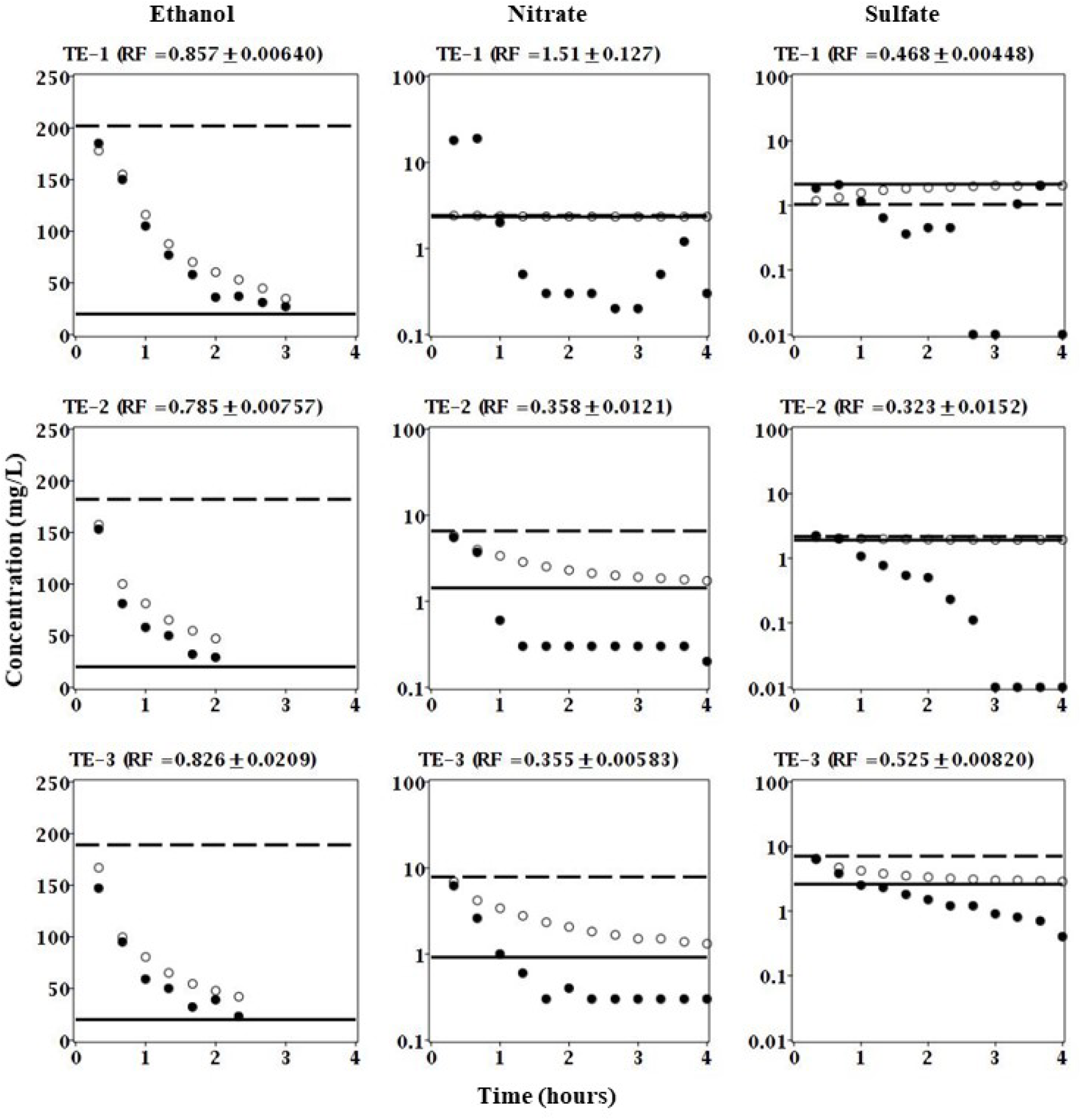
Breakthrough curves of ethanol, nitrate, and sulfate for treatment exposures 1 through 6 (TE-1 through TE-6) in well FW222, solid circles (●) are measured concentrations in the extraction fluid, open circles (○) are expected concentrations in the extraction fluid based on bromide (non-reactive tracer), dashed line (□ □) is the concentration in the injection fluid, solid line (□ □) is the concentration in the aquifer fluid or the lower detection limit.

The rate of groundwater flow was so high for exposures four, five, and six (Fig. 3) that the concentration of ethanol was diluted to below the method detection limit (20 mg/L) within the first hour and therefore only two or three data points were available for analysis (data not shown). Acetate production was observed for exposures one, two, and three as evident by recovery factors greater than one (data not shown). However, given that acetate is an intermediate byproduct of ethanol reduction and can serve as an electron donor for further reduction its temporal behavior is somewhat difficult to interpret beyond evidence of ethanol oxidation.

The breakthrough curves of ethanol, nitrate, and sulfate for exposure seven in the treatment well (TE-7) demonstrated concomitant ethanol and nitrate removal as evident by substantially lower than expected concentrations; again, nitrate concentrations actually fell below even that of the aquifer fluid (Fig. 6). Moreover, the recovery factors for both ethanol and nitrate were much less than one, 0.796 and 0.789, respectively. In contrast, the breakthrough curves for exposure one in the control well (CE-1) were similar to exposure one in the treatment well (TE-1) which did not demonstrate concomitant ethanol and nitrate removal; nitrate concentrations in the control well (CE-1) were nearly identical to those expected due to dilution (Fig. 6). Moreover, the recovery factor for nitrate was nearly equal to one, 0.952 to be exact (Fig. 6). Interestingly, the recovery factor for ethanol was less than one, 0.865 to be exact (Fig. 6). Moreover, acetate production was also observed, although substantially less as compared to the treatment well (data not shown). One explanation for the apparent removal of ethanol but not nitrate in the first exposure of the control well (Fig. 6) and the first exposure of the treatment well (Fig. 5) is the presence of oxygen as a higher energy yielding electron acceptor. For example, it is likely that oxygen was introduced to the injection fluid during the above ground mixing of bromide and ethanol. Therefore, it is likely that aerobic respiration of ethanol occurred rapidly and prior to the onset of anaerobic conditions where nitrate would be the next highest energy yielding electron acceptor.

**Fig. 6.**
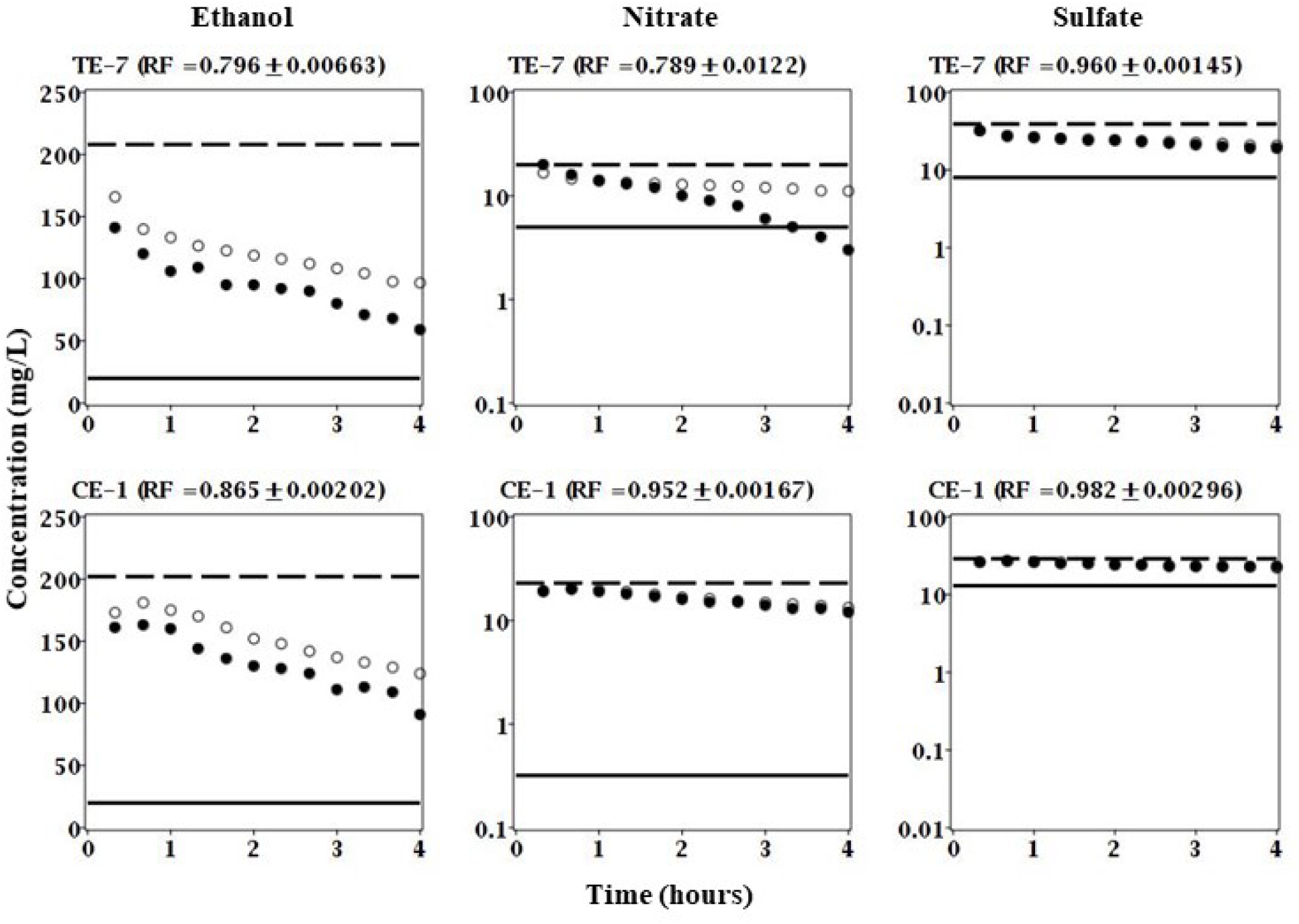
Breakthrough curves of ethanol, nitrate, and sulfate for treatment exposure 7 (TE-7) in well FW222 and control exposure 1 (CE-1) in well FW224; both occurring in week 14 (Table 1), solid circles (●) are measured concentrations in the extraction fluid, open circles (○) are expected concentrations in the extraction fluid based on bromide (non-reactive tracer), dashed line (□ □) is the concentration in the injection fluid, solid line (□ □) is the concentration in the aquifer fluid or the lower detection limit.

Overall, these results strongly suggested that the treatment well sustained its ability for nitrate removal even in the absence of ethanol for up to six weeks. It is conceivable that the duration of sustained ability could have lasted much longer and therefore additional *in situ* studies are needed to constrain an upper limit on the duration of this phenomenon.

### 3.2. Microbial Community Structure

NMDS was conducted to assess the similarity of the natural microbial communities at the level of OTU (Fig. 7). The number of dimensions was increased from two to three at which the ordination stress decreased from approximately 4 to 0.5 and remained below 0.5 up to at least seven dimensions (scree plot not shown). The NMDS plots showed that the control well clustered more closely as compared to treatment well (Fig. 7). These results suggested that exposure to ethanol caused a notable shift in the microbial community as compared to no exposure to ethanol. The microbial community in the control well at week four (W04) and after exposure to ethanol at week 14 (W14*) were notably dissimilar to the other time points (Fig. 7). These results suggested that the microbial community shifted in response to no added electron donor (W04) and added electron donor (W14*) conditions. However, the microbial communities in both the control and treatment wells were notably similar at weeks 14 (W14) and one (W01) (Fig. 7). These results suggested that by week 14 (W14) both microbial communities shifted back to a structure that was notably similar to their initial condition at week one (W01). These results were particularly surprising when considering that the treatment was exposed to six consecutive weeks of ethanol whereas the control was not.

**Fig. 7.**
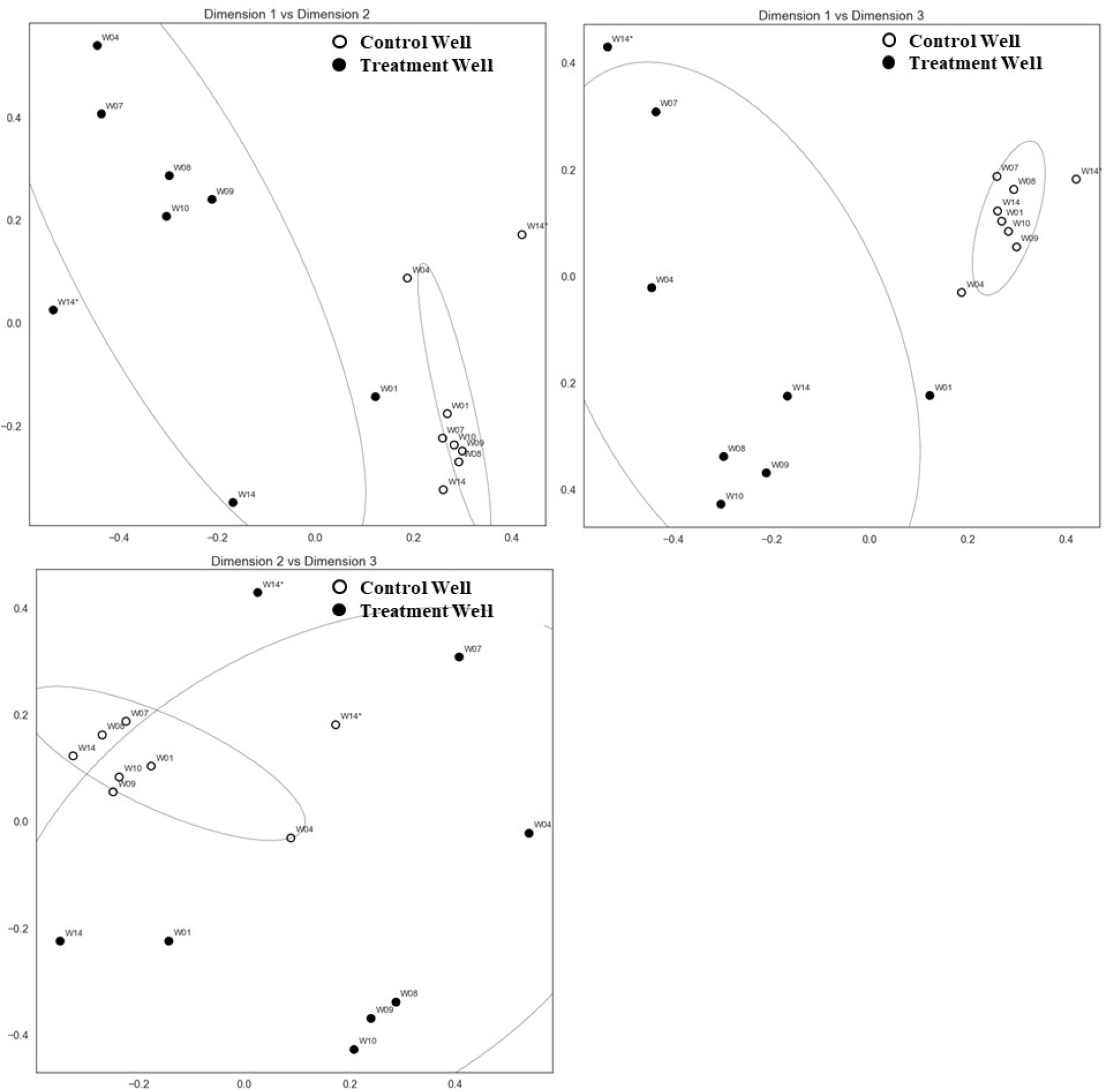
Non-metric multi-dimensional scaling (NMDS) plots during the 14-week experiment (W01 through W14*) for the control and treatment wells, *indicates post-ethanol exposure.

HCA was conducted to further assess the similarity of the natural microbial communities at the level of OTU (Fig. 8). The communities clustered into four distinct groups (G1 through G4) (Fig. 8). Group 1 consisted entirely of the control well whereas groups 2, 3, and 4 consisted entirely of the treatment well (Fig. 8). Within the control well (G1), the community after exposure to ethanol (W14*) was most dissimilar as indicated by the dendrogram (Fig. 8). This result was expected based on the NMDS plots (Fig. 7). Group 2 consisted of the treatment well at weeks one (W01) and the beginning of week 14 (W14), which were more similar to each other than to any other time points across both exposure treatment and exposure control (Fig. 8). This was also consistent with the NMDS results (Fig. 7). The HCA quantified the similarity as 0.67 on a scale of zero being most similar and one being least similar (Fig. 8). Therefore, both the NMDS and the HCA suggested that the microbial community in the treatment well did not sustain its ability in response to exposure to ethanol (Figs. 7 and 8). This was particularly surprising when considering that the breakthrough curves in the treatment well strongly suggested that the community sustained its ability for ethanol-induced nitrate removal (Figs. 5 and 6). Group 3 consisted of the treatment well at weeks eight, nine, and ten whereas group 4 consisted of weeks four, seven, and week 14* (Fig. 8). In terms of timing with respect to ethanol exposure, group 3 coincided with the six-week period of no exposure to ethanol whereas group 4 coincided with the initial and final exposure to ethanol (Fig. 8 and Table 1). These results were expected based on the timing of ethanol exposures. As previously mentioned, the most surprising result was the relatively high similarity of the community structures of the treatment well at week one (W01) and the beginning of week 14 (W14) (Figs. 7 and 8) despite the apparent sustained ability for ethanol-induced nitrate removal (Figs. 5 and 6). It is possible that the sessile microbial community was readily able to utilize ethanol but without sediment samples this could not be tested. It is also possible that genetic alterations, rather than persistent changes to the community structure, were the primary mechanism that allowed the exposure treatment to respond rapidly to ethanol exposure (W14*).

**Fig. 8.**
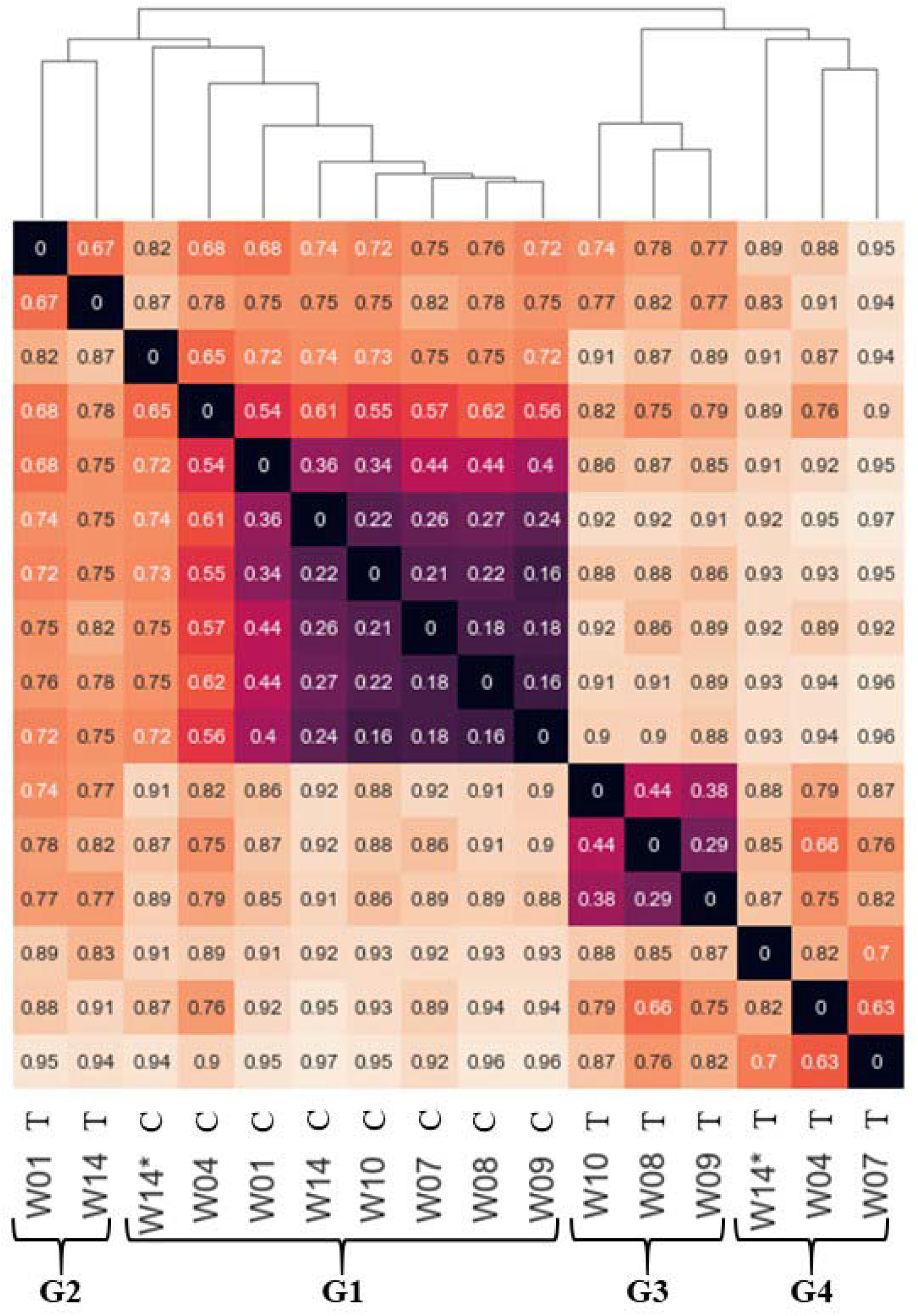
Hierarchical clustering analysis of operational taxonomic units (OTUs) during the 14-week experiment (W01 through W14*) for the control (C) and treatment (T) wells, *indicates post-ethanol exposure, G1, G2, G3, and G4 indicate distinct groupings.

Relative abundance analysis was conducted to assess the shifts in particular taxa at the level of phylum (Fig. 9). The microbial community of the control well was dominated by *Proteobacteria* for weeks one through the beginning of 14 but showed considerable variability (Fig. 9). The relative abundance of other taxa in the control well, such as *Nitrospirae, Firmicutes*, and *Woesearchaeota* were also notable for weeks one through the beginning of 14 and showed considerable variability (Fig. 9). During this time, the control well was not exposed to ethanol (Table 1). Therefore, the temporal changes in taxa in the control well for weeks one through 14 were representative of natural biogeochemical conditions. The high relative abundance and temporal variability of *Proteobacteria, Nitrospirae*, and *Firmicutes* under natural biogeochemical conditions was expected based on a recent study at the ORR by King et al. (2017). King et al. (2017) demonstrated similar results from *in situ* above ground bioreactors and noted that such taxa are associated with low dissolved oxygen and/or representative of nitrate reducers. Both low dissolved oxygen and the presence of nitrate are characteristic of the dissolved-phase chemistry at the study site (Paradis et al. 2016). The control well was exposed to ethanol during the middle of week 14 (W14) and sampled for microbial community structure at the end of week 14 (W14*) (Table 1). After exposure to ethanol (W14*), *Acidobacteria* substantially increased in relative abundance, replacing *Proteobacteria* as the dominant phylum (Fig. 9). These results differ from previous studies at the ORR which showed increases of *Proteobacteria* and decreases of *Acidobacteria* after exposure to ethanol (Spain et al. 2007; Cardenas et al. 2008). However, those studies characterized the microbial communities associated with sediment (sessile) and after prolonged (three weeks to two years) exposures of ethanol (Spain et al. 2007; Cardenas et al. 2008) whereas this study characterized microbial communities associated with groundwater (planktonic) and after a brief (less than four hours) exposure of ethanol. It is possible that the sessile microbial community changed in a manner consistent with previous studies, but this is not known due to lack of sediment samples. It is also possible that duration of exposure to ethanol, i.e., prolonged versus brief, had a notable effect on the relative abundance of taxa as previously demonstrated by Pernthaler et al. (2001). Nevertheless, these results demonstrated that the planktonic microbial community in the control well was relatively stable under natural conditions but rapidly changed after exposure to ethanol.

**Fig. 9.**
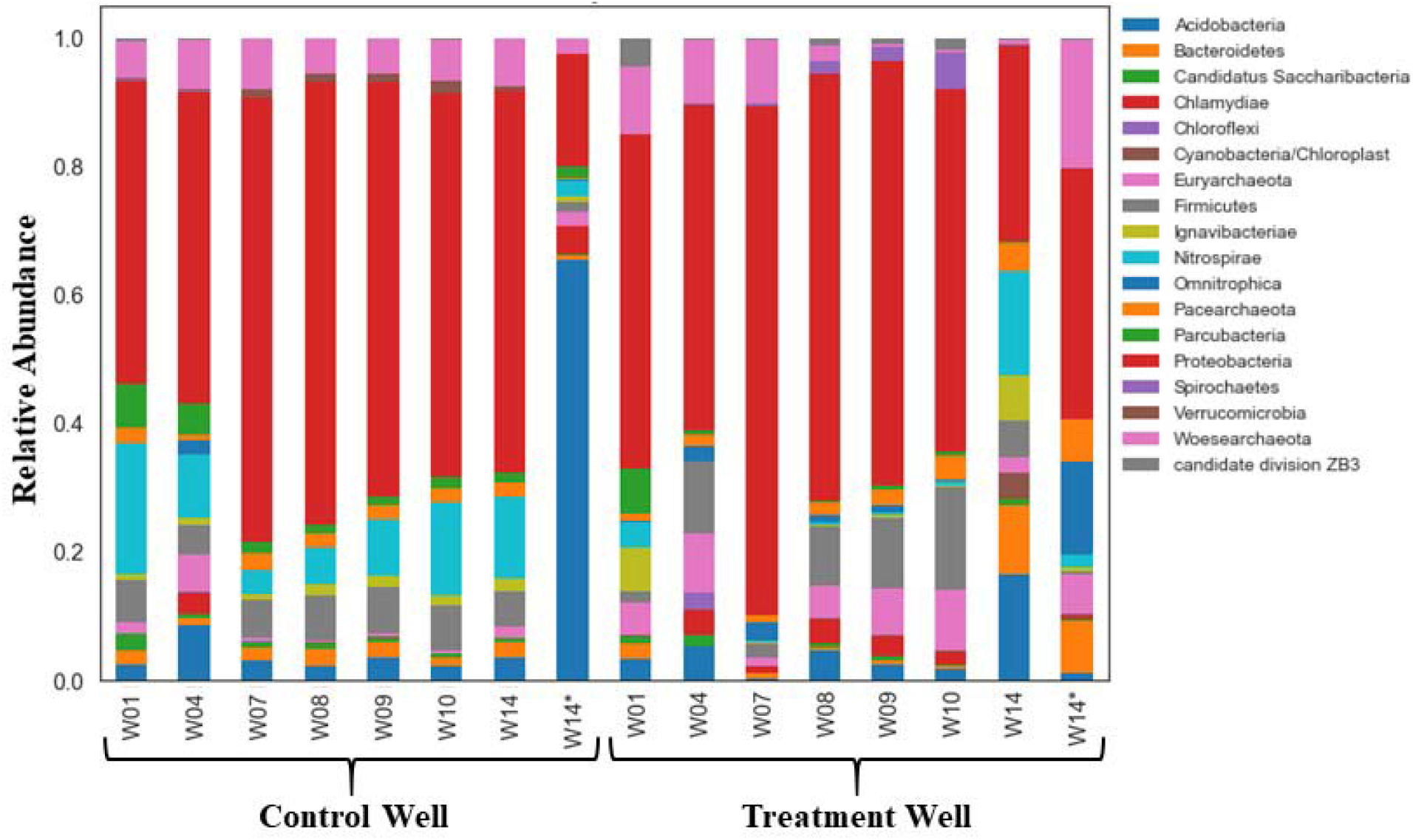
Relative abundance of microbial taxa at the phylum level during the 14-week experiment (W01 through W14) for the control and treatment wells, *indicates post-ethanol exposure.

The treatment well was dominated by *Proteobacteria* for weeks one through 10 but varied considerably more than the control well (Fig. 9). The relative abundance of other taxa in the treatment well, such as *Firmicutes* and *Woesearchaeota* were also notable for weeks one through 10 and showed considerable variability (Fig. 9). Compared to the control well during this time, the community in the treatment well by week 10 was notably different than week one (Fig. 9). A notable change in the community in the treatment well was expected because by week 10 the treatment had been exposed to six consecutive weeks of ethanol whereas the exposure control had not been exposed to ethanol (Table 1). By the beginning of week 14, the treatment well had been exposed to ethanol for six consecutive weeks followed by six consecutive weeks without exposure to ethanol (Table 1). As compared to the control well, the community in the treatment well by week 14 was notably different than week one (Fig. 9). Therefore, if the microbial community in the treatment was able, and sustained its ability for, ethanol-induced removal of nitrate, which the breakthrough curves strongly suggested (Fig. 6), then the community at the beginning of week 14 (W14) may be representative of a sustained community (Fig. 9). Likewise, if the microbial community in the control well lacked the sustained ability for ethanol-induced removal of nitrate, which the breakthrough curves strongly suggested (Fig. 6), then the community at the beginning of week 14 (W14) may be representative of a non-able community (Fig. 9). The relative abundance of taxa in the treatment well after its final exposure to ethanol (W14*) was notably different than before its final exposure to ethanol (W14) as indicated by the increase of *Woesearchaeota* and decrease of *Nitrospirae* (Fig. 9). These results demonstrated that the microbial community in the exposure treatment changed upon exposure to ethanol and sustained a level of ability in the absence of exposure to ethanol. As previously noted, it is also possible that genetic changes, rather than persistent changes to the community structure, were the primary mechanism that allowed the treatment well to respond rapidly to ethanol exposure (W14*). Therefore, future *in situ* studies of sustained ability should attempt to characterize the sessile community as well as investigate the genetic changes to ethanol exposure.

## 4. Conclusions

The objectives of this study were to establish a natural microbial community able to remove nitrate from groundwater via the addition of an electron donor and then determine how long this ability could be sustained in the absence of the electron donor and elucidate the microbial mechanism(s) responsible for this ability. The results of this study strongly suggested that the *in situ* ability of a natural microbial community to remove nitrate from groundwater can be sustained in the prolonged absence of an electron donor; in this case, at least six weeks in the absence of ethanol. Moreover, this ability was not be revealed in the experiment by a sustained and selected enrichment of a planktonic microbial community based on 16S rDNA. However, it is possible that such a microbial community may be present in the sessile state or that the predominant mechanism(s) of this ability exist at the enzymatic- and/or genetic-levels. Nevertheless, this study demonstrated that the exposure history of groundwater to an electron donor can play an important role in the removal of nitrate.

## Acknowledgements

This material by the ENIGMA-Ecosystems and Networks Integrated with Genes and Molecular Assemblies (http://enigma.lbl.gov), a Science Focus Area Program at Lawrence Berkeley National Laboratory is based upon work supported by the U.S. Department of Energy, Office of Science, Office of Biological & Environmental Research under contract number DE-AC02-05CH11231. Oak Ridge National Laboratory is managed by UTBattelle, LLC, for the U.S. Department of Energy under contract DE-AC05-00OR22725.

